# The SARS-CoV-2 Y453F mink variant displays a striking increase in ACE-2 affinity but does not challenge antibody neutralization

**DOI:** 10.1101/2021.01.29.428834

**Authors:** Rafael Bayarri-Olmos, Anne Rosbjerg, Laust Bruun Johnsen, Charlotte Helgstrand, Theresa Bak-Thomsen, Peter Garred, Mikkel-Ole Skjoedt

**Author notes:** **Corresponding author:** Mikkel-Ole Skjødt.

## Abstract

SARS-CoV-2 transmission from humans to animals has been reported for many domesticated species, including cats, dogs and minks. Identification of novel spike gene mutations appearing in minks has raised major concerns about potential immune evasion and challenges for the global vaccine strategy. The genetic variant, known as “cluster-five”, arose among farmed minks in Denmark and resulted in a complete shutdown of the world’s largest mink production. However, the functional properties of this new variant are not established. Here we present functional data on the Y453F cluster-five receptor-binding domain (RBD) and show that it does not decrease established humoral immunity or affect the neutralizing response in a vaccine model based on wild-type RBD or spike. However, it binds the human ACE-2 receptor with a four-fold higher affinity suggesting an enhanced transmission capacity and a possible challenge for viral control.

## Introduction

Experimental infection models have shown that SARS-CoV-2 is capable of infecting a wide range of animal species, such as cats, dogs, ferrets, and hamsters (*1–3*), and natural reverse-zoonotic transmission to animals has been documented in farmed mink (*4*), dogs (*5, 6*), and felines (*6–8*). Several variants affecting the spike protein have been documented since the early reports in the spring of 2020 of SARS-CoV-2 transmission from humans to minks (*4, 9, 10*). The spike protein mediates the binding of SARS-CoV-2 to the human angiotensin-converting enzyme-2 (ACE-2) receptor (*11, 12*), and as such, determines its infectivity and transmissibility. Moreover, the spike protein is the main target for the host’s protective antibody response (*11, 13, 14*). Mutations arising through antigenic drift may thus result in novel viral escape strains. One of these cluster variants, disclosed as “cluster-five”, has been discovered in Denmark and bears a tyrosine to phenylalanine substitution (Y453F) in the receptor-binding domain (RBD) of the spike protein (*15*). This position is conserved between SARS-CoV-1 and −2, and analyses of the crystal structure of SARS-CoV-2 RBD in complex with ACE-2 reveal that it is directly involved in the interaction of the virus with the host cell receptor (*16, 17*). However, the role of this variant concerning functional transmission and establishment of immunity is unknown. At present two new RBD variants known as N439K and N501Y have been found in humans. The latter appears to have a better transmission or improved immune evasion than the original counterpart and this variant is currently increasing in the infection cases in the UK (*18*).

There is a general lack of evidence characterizing the pathogenesis, functional properties, and impact on these new cluster variants’ immune recognition and memory responses. Despite this, there have been speculations and reports suggesting that the cluster-five variant might have evasion potentials by reducing the immunity of convalescent individuals and that it could even challenge the global vaccine strategies that are being initiated currently (*15*). Based on these presumptions, the Danish government in November 2020 decided to completely shut down all Danish mink farms, representing the largest mink fur production in the world.

To address the biophysical characteristics and the impact of this variant on established immunity, we expressed recombinant SARS-CoV-2 RBD “wild-type” (wt, from the Wuhan-Hu-1/2019 isolate) and “cluster-five” Y453F and the ACE-2 ectodomain. The Y453F variant did not alter the neutralization potency of sera from convalescent individuals exposed to the wt strain in the spring of 2020, nor did it challenge the humoral neutralizing vaccine response in a mouse model immunized with wt RBD or prefusion-stabilized spike protein ectodomain. However, we found a four-fold affinity increase of the Y453F variant to the ACE-2 binding that could result in an enhanced spreading potential.

## Results

### Biophysical characterization

To determine the biological significance of the Y453F mutation, we studied the impact on protein stability and function by thermal denaturation and binding kinetics experiments (**Fig. 1**). Using the ratio of the intrinsic fluorescence at 350 and 330 nm over a temperature gradient from 35 to 95 °C, we observed no significant differences in the inflection temperatures (Ti) (53.43 and 53.37 °C for the wt RBD and Y453F, respectively), suggesting that the variant has no critical effect on protein stability (**Fig. 1B**). We also determined the kinetics of the interaction with the ectodomain of human ACE-2 by biolayer interferometry (**Fig. 1C, D**). The Y453F variant bound with a four-fold higher affinity than the wt (3.85 nM vs 15.5 nM), and analyses of the Ka and Kdis revealed that it bound faster to ACE-2 (Ka 1.5 x 10^5^ Ms vs 4 x 10^5^ Ms) and remained bound for longer (7 x 10^−4^ s^−1^ vs 6 x 10^−3^ s^−1^).

**Figure 1.**
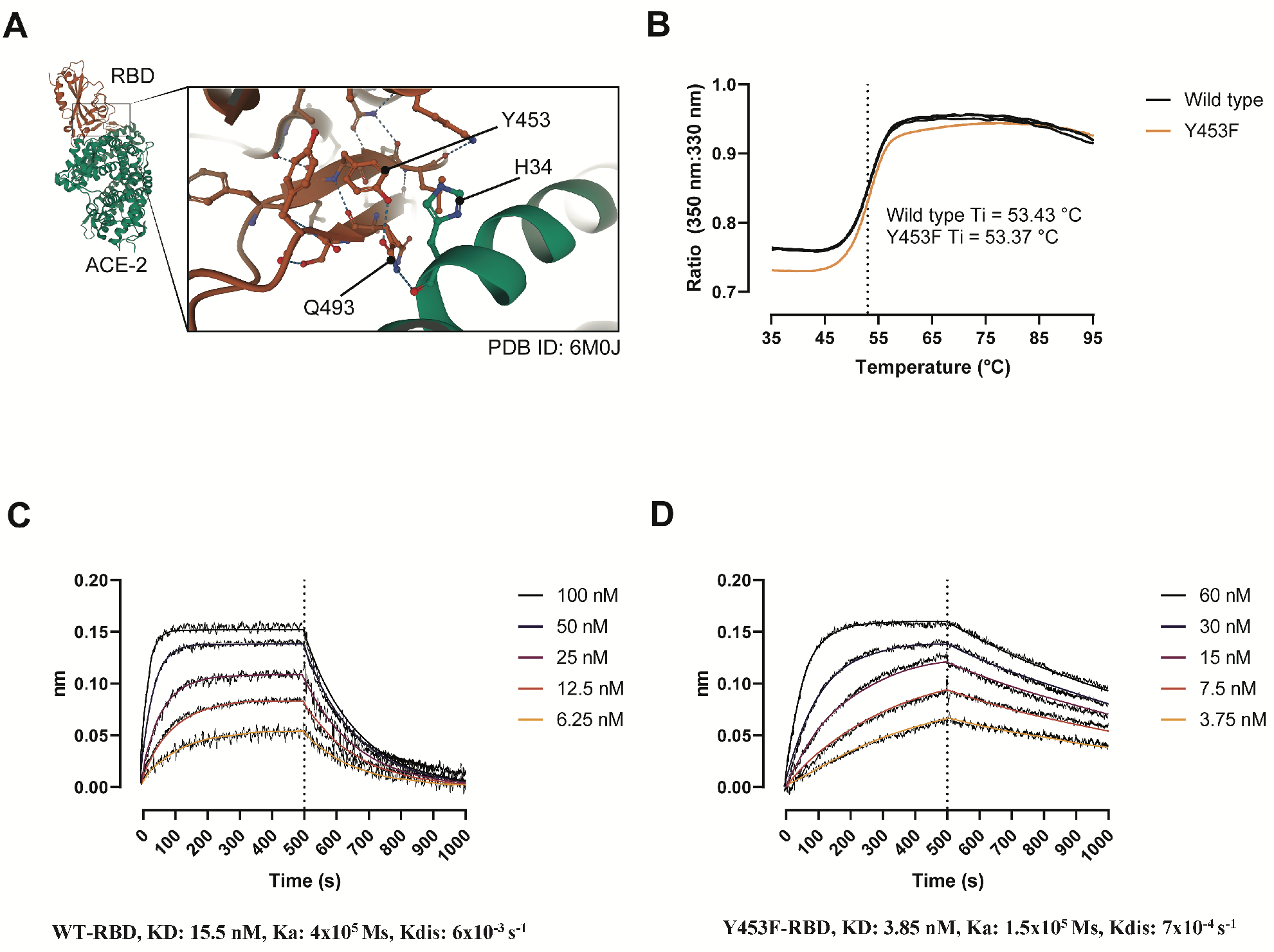
**(A)** Details of the binding interface of SARS-COV-2 RBD (orange) and the human ACE-2 receptor (green). The residue 453 maps to the receptor-binding motif within the RBD (residue 438–506) and directly engages the N-terminal helix of the peptidase domain of ACE-2 (*19*). Dashed lines represent non-covalent interactions. Created with PDBe Molstar (https://molstar.org/). **(B)** Thermal denaturation curves of the wt and Y453F RBD. Data is represented as the 350:330 nm ratio of each replicate. Ti, inflection temperature. **(C, D)** wt (C) and Y453F (D) RBD binding response curves to ACE-2 determined by biolayer interferometry. ACE-2-Fc was immobilized unto anti-human IgG Fc capture sensors and dipped into serial dilutions of RBD (5-point 2-fold dilutions starting at 100 nM [wt] or 60 nM [Y453F]).

### Determination of the neutralization capacity of COVID-19 convalescent patients against the Y453F variant

To clarify whether the tyrosine to phenylalanine substitution in the surface of the RBD affects not only its functionality but also recognition by the immune system, we determined the neutralization capacity of sera from qPCR-confirmed COVID-19 convalescent individuals (*n* = 141) using a previously described antibody neutralization ELISA (*20*). We observed a tight linear relationship (*R^2^* = 0.9430) and a highly significant correlation (*ρ* = 0.9711, *p* < 0.0001) of the serum antibody neutralization capacity to both RBD variants (**Fig. 2**).

**Figure 2.**
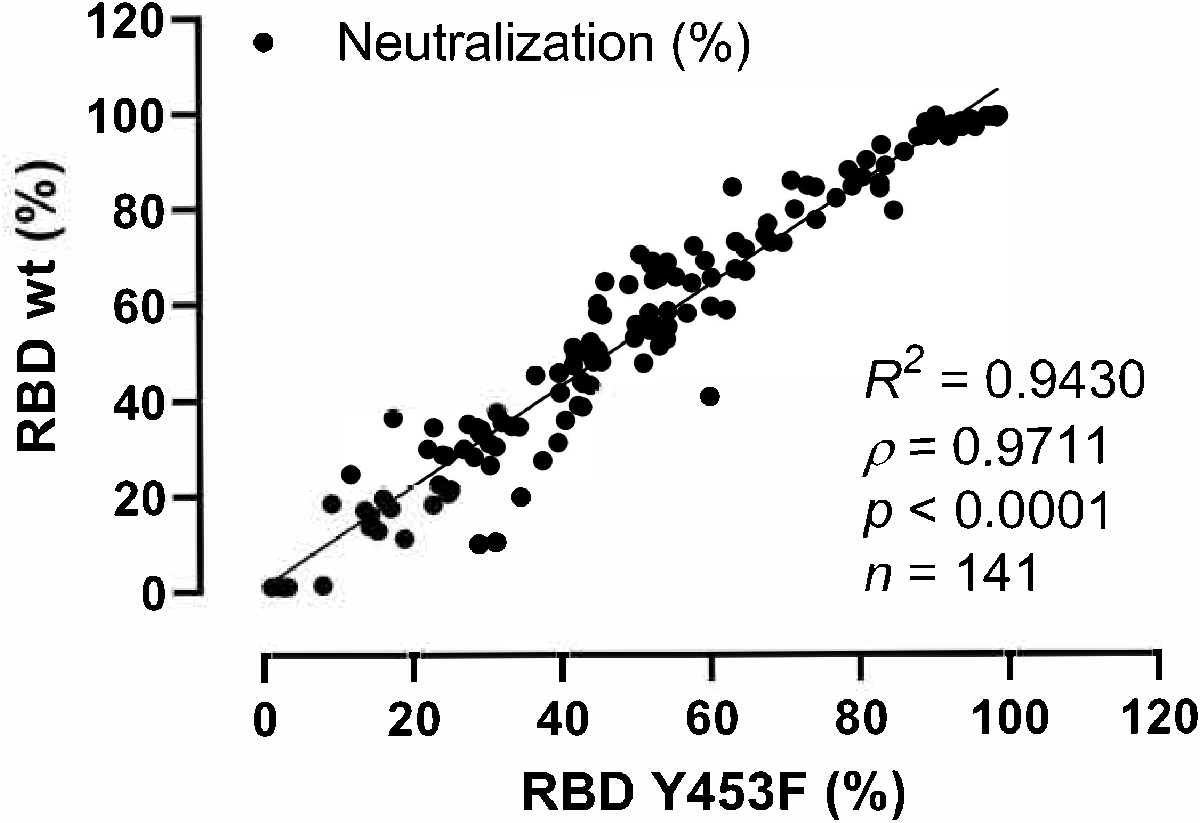
Neutralization potency of COVID-19 convalescent patient sera / Sera from COVID-19 patients neutralization of the wt and Y453F RBD. Linear regression and Spearman rank correlation analyses of the neutralization potency of COVID-19 patient sera against wt and Y453F RBD. Trend line represents a linear regression. Negative neutralization values were normalized to 0.

### Determination of the vaccine response in a pre-clinical animal model

Next, we evaluated the neutralization capacity of polyclonal sera and mouse monoclonal antibodies (mAbs) isolated from mice immunized with wt RBD or prefusion-stabilized spike protein as a pre-clinical vaccine model (described elsewhere, (*20*)) (**Fig. 3**). Sera from RBD immunized mice neutralized the wt and Y453F RBD with an IC_50_ of 28,369 and 26,579, respectively, several fold more effectively than sera from spike immunized mice (IC_50_ of 7,075 and 6,363, respectively) (**Fig. 3A**). More importantly, and regardless of the immunization strategy, the mice sera did not show differences in the neutralization potency of the two RBD variants.

**Figure 3.**
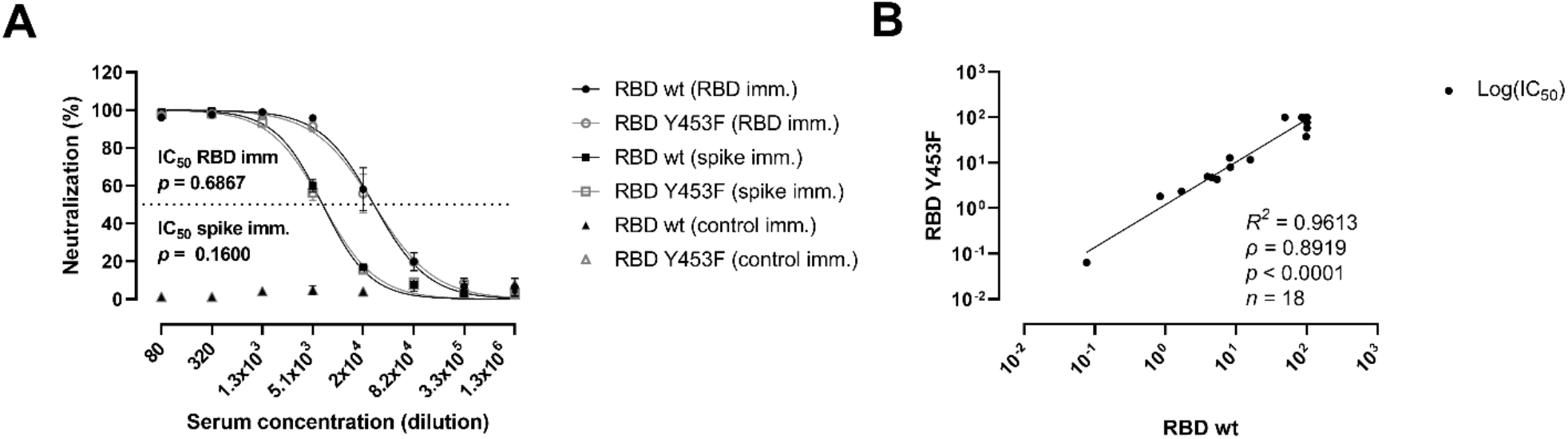
Neutralization potency of polyclonal sera and mAbs from mice immunized against SARS-CoV-2 wt spike or RBD. **(A)** Comparison of the best-fit IC_50_ of polyclonal sera from mice immunized with SARS-CoV-2 wt RBD (RBD imm.) or spike (spike imm.) against wt RBD (black symbols) and the Y453F variant (grey symbols). A SARS-CoV-2 nonrelated immunization (control imm.) was used as control. Connecting lines represent the nonlinear fit. Data are presented as mean ± SEM. **(B)** Linear regression and Spearman rank correlation analyses of the neutralization potency (log[IC_50_]) of the wt and Y453F RBD using individual mAbs raised against wt RBD (*n* = 10) or wt spike (*n* = 8).

Finally, we compared the neutralization potency of high-affinity mAbs (*n* = 18) derived from immunized mice (**Fig. 3B**). Linear regression and Spearman rank correlation analyses of the IC_50_ values against wt and Y453F RBD revealed that the neutralization potencies against both variants correlated strongly (*R^2^* = 0.9613, *ρ* = 0.8919, *p* < 0.0001). The neutralization potential of the individual mAbs neutralization potential was assessed separately.

## Discussion

The emergence of nonsynonymous mutations in the SARS-CoV-2 spike gene has been reported since the beginning of 2020. The main part of these mutations have been identified within the European continent (>20), with fewer identifications on the Asian and American continents (*21*). The vast majority of these residue substitutions are located in the spike regions outside of the receptor-binding domain (RBD) with the D614G mutation being a common variant reported on all continents and which seems to be taking over likely due to selective advantages (*22–26*). Recently, three new genetic mutations inducing residue changes in the RBD have been reported in Europe, i.e. N439K, Y453F, and N501Y. The N439K may make the virus more infectious due to an increased affinity towards ACE-2 and/or a reduced sensitivity to neutralizing antibodies (*26–28*). The N501Y mutant is a part of a novel strain, “Variant of Concern 202012/01”/B.1.1.7, that has accumulated 17 mutations—8 of them in the spike gene—and in some areas of England may account for most of the new cases at the time of writing (*18*). The Y453F was first identified in Denmark in the summer of 2020 among farmed minks. The variant was coupled to the “cluster five” mutation (*15*), and arose a great international concern when it was reported to have been transmitted to humans. Millions of minks were culled and their pelts discarded, effectively shutting down a country-wide industry. However, a thorough functional characterization of the impact of the Y453F RBD variant on immunity and ACE-2 interaction has so far not been conducted.

When we challenged the RBD:ACE-2 interaction with COVID-19 convalescent sera from 141 qPCR-confirmed individuals that have been infected with the original SARS-CoV-2 variant, we found no reduction in the serum capacity to neutralize the Y453F variant. This conflicts with a preliminary report that found plasma samples from convalescent individuals with low and intermediate neutralization titers had a reduced neutralization potency against the variant (*15*). To further address this, we used a traditional vaccine approach applying mice immunized with wt RBD or the full wt ecto-domain spike and assessed the antibody titers and neutralization capacity of wt and Y453F RBD. In the vaccine models, we found no difference concerning inhibition of the RBD/ACE-2 interaction of the two variants, which was also the case when we challenged 18 different neutralizing monoclonal antibodies raised against wt RBD or spike antigens.

The tyrosine at position 453 has been shown to be a critical residue engaged in direct interaction with the ACE-2 receptor via the tyr OH group (*16*). We therefore expected that there would be either a reduced or similar affinity with the substitution to a phenylalanine. However, to our surprise we found a four-fold higher affinity of the 453F RBD binding to ACE-2 (KD 3.85 nM) compared to wt RBD. The precise reason for this remains to be established but there are several other tyrosine reesidues in RBD that might be utilized in the ACE-2 interaction. The concequence of this affinity increase on the physiological avidity could even be more pronounced for the trimeric spike/ACE-2 interaction.

Regardless, this suggests two essential points with the developing clusters of spike variants: That immunity might not be dramatically challenged and that emphasis should also be focused on characterizing the transmission capacity and interaction properties of new emerging SARS-CoV-2 strains. This follows in the line of the new UK discovered N501Y mutant that within short time gained transmission dominance in many areas including London, where by mid-December represented more than 60% of the cases (*29*). Functional properties on the N501Y mutant variant is not yet established.

Taken together, the results suggest that the molecular evolution of the SARS-CoV-2 virus has sofar been taken it in the direction of receptor affinity adaptation and optimization rather than towards an immune evasion strategy.

## Data availability

All data are contained within this manuscript.

## Acknowledgements

The authors would like to thank Jais Rose Bjelke from Novo Nordisk A/S for quality analysis of the recombinant proteins and Mss Camilla Xenia Holtermann Jahn, Sif Kaas Nielsen, and Jytte Bryde Clausen, from the Laboratory of Molecular Medicine for their excellent technical assistance.

## Authors’ contributions

RB-O and MOS conceived and designed the study; RB-O, and CH enabled recombinant protein production; RB-O, AR, LBJ and MOS performed experiments; RB-O, LBJ and MOS analyzed the data; RB-O, PG and MOS wrote the paper with inputs from all co-authors. All authors approved the final version of the manuscript.

## Funding and additional information

This work was financially supported grants from the Carlsberg Foundation (CF20-0045 - MOS, RBO & PG) and the Novo Nordisk Foundation (NFF205A0063505 and NNF20SA0064201 - MOS, RBO & PG).

## Abbreviations

RBD: Receptor-binding domain
ACE-2: Angiotensin-converting enzyme 2
BLI: Biolayer interferometry

